# Nucleic acid strand length governs mitochondrial reprogramming and mtROS-associated antiviral responses following TLR3 engagement

**DOI:** 10.64898/2025.12.03.691914

**Authors:** Duale Ahmed, Malak Al Daraawi, Allan Humphrey, Omar Abdo, David Roy, Mary-Elizabeth Sheridan, Zoya Versey, Rayhane Mejlaoui, Allison Jaworski, Alex Edwards, Alfonso Abizaid, Ashok Kumar, Ashkan Golshani, Edana Cassol

## Abstract

Mitochondria are important rheostats that regulate innate sensing processes by producing energy, biosynthetic precursors, and bioactive molecules that affect cellular signaling. When viral nucleic acids engage endosomal and cytosolic pattern recognition receptors (PRR), antiviral immune responses are supported by mitochondrial remodeling but the role of mitochondria in fine tuning ligand-specific responses remains incompletely understood. For example, endosomal TLR3 can detect various lengths of dsRNA (0.4-8 kb) ranging from viral segmented genomes or endogenous nucleic acids have been shown to induce distinct cytokine profiles. However, it is unclear if these differences are associated with differential mitochondrial remodeling. Here, we report that TLR3 engagement with both high (HMW; 1.5-8 kb) or low molecular weight (LMW; 0.2-1 kb) Polyinosinic:polycytidylic acid (Poly(I:C)) is associated with reduced but sustained oxidative phosphorylation (OXPHOS) activity and increased mitochondrial reactive oxygen species (mtROS) production/accumulation to support antiviral responses in bone marrow-derived macrophages (BMDM). They differed in the amount of mtROS production, their spare respiratory capacity (SRC) and their mitochondrial membrane potential (MMP). Interestingly, while uncoupling protein 2 (UCP2) was found required for antiviral cytokine production, it did not contribute to ligand specific responses. Dynamic modulation of complex I of the electron transport chain (ETC), however resulted in the differential accumulation of mtROS (HMW>LMW). Further, selectively targeting the mtROS derived from Complex I leads to augmented type I IFN production. Overall, these findings highlight that targeting specific sources of mtROS without affecting electron flow may be a potential avenue for specific augmentation of antiviral responses during viral infections.

## Introduction

Macrophages are tissue resident cells whose function is highly influenced by their local microenvironment^1–8^. These cells play a central role in initiating responses to injury and infection by regulating inflammatory processes and immune cell infiltration^1,2^. At resolution, they also remove cellular debris and support tissue remodeling and repair^3,4^. These responses are initiated by pattern-recognition receptors (PRRs), which detect pathogen-associated (PAMPs) and/or damage-associated molecular pattern molecules (DAMPs)^9–13^. In viral infections, this includes nucleic acids, which interact with endosomal and cytosolic PRR to induce type I interferon (IFN) and inflammatory cytokine production and associated responses^11–13^. To date, we have a limited understanding of how innate sensing of viruses differ across cell and tissue types and how they are modified by the local microenvironment including oxygen and nutrient availability, which alter the cell’s metabolic status^14–19^.

Mitochondria are increasingly recognized as a rheostat that fine-tunes macrophage activation and function^16,19–24^. Beyond supporting bioenergetic and biosynthetic demands of the cell by metabolizing glucose^15,25–28^, glutamine^29,30^, branched-chain amino acids^31–33^, and fatty acids^34,35^, mitochondria generate intermediate metabolites and bioactive molecules (e.g., tricarboxylic acid cycle metabolites, ROS) that regulate cellular signaling, transcription and epigenetics^16,18,21,36–40^. They also modulate immune activation through processes such as OXPHOS^16,19,41^, mitochondrial fusion/fission dynamics^42–45^, and functioning as a scaffold to support and regulate proper activation^23,46–50^. Synthetic nucleic acids that engage PRR (e.g., Polyinosinic:polycytidylic acid (Poly(I:C)), Imiquimod (R837), and CpG oligodeoxynucleotides (ODNs) have been shown to require some degree of mitochondrial remodeling to elicit responses^18,51–55^. However, since standard cell culture conditions include supraphysiological levels of most nutrients, such as glucose^56^. This may alter mitochondrial reprogramming *in vitro*^57^, obscuring aspects of mitochondrial function that contribute to the fine tuning of ligand specific responses.

Toll-like receptor 3 (TLR3) offers a unique opportunity to investigate how mitochondria contribute to the fine-tuning of ligand specific responses. This endosomal receptor detects a range of lengths of dsRNA (min: 40-50bp; max: >8kb), including diverse self-derived dsRNA or dsRNA viruses with segmented genomes^58–63^. Studies have shown that short and long dsRNA differentially engage TLR3 to induce distinct activation states across cell types^64–66^. Specifically, human monocyte-derived dendritic cells and mouse RAW264.7 cells have been found to be more responsive to shorter dsRNAs, whereas mouse embryonic fibroblasts produce stronger inflammatory/antiviral responses to longer dsRNAs^64,65^. We found that TLR3 engagement by HMW (1.5-8 kb) vs. LMW (0.2-1 kb) Poly(I:C) differentially induces NF-κB-driven inflammation in a HIF-1α-dependent manner^66^. Further, while HMW Poly(I:C) requires both OXPHOS activity and Complex III-derived mtROS to support antiviral signaling^18^, it is unclear whether LMW Poly(I:C) uses the same mechanisms.

In this study, we investigated whether HMW vs. LMW Poly(I:C) differentially remodel mitochondrial function to regulate type I IFN responses in murine BMDMs. We found that both forms of Poly(I:C) reduced OXPHOS activity, but that HMW induced a more pronounced loss of spare respiratory capacity (SRC%) and mitochondrial membrane potential (MMP), independent of ligand concentration or potency. While the inner mitochondrial protein uncoupling protein 2 (UPC2) supported type I IFN production, it did not contribute to ligand differences. Instead, altered Complex I activity primarily drove the divergent mitochondrial function between HMW and LMW activation. In fact, preventing Complex I-derived mtROS without impairing OXPHOS activity resulted in differential effects on type I IFN secretion, suggesting ETC flux may function as a rheostat for regulating ligand-specific antiviral responses through the same PRR.

## Materials and Methods

### Reagents

High glucose Dulbecco’s Modified Eagle Media (DMEM) with 4 mM L-glutamine and 1 mM sodium pyruvate (11995-073), glucose/glutamine/pyruvate/phenol red-free DMEM (A14430-01), fetal bovine serum (FBS) (10437-028), penicillin/streptomycin (PenStrep) (15140-122), glucose (A2494001), sodium pyruvate (11360-070), L-glutamine (25030-081), Tetramethylrhodamine, methyl ester (TMRM) (I34361), MitoTracker^TM^ Green (M7514), MitoSOX^TM^ Red (M36008), Propidium iodide (PI) (P1304MP) as well as antibodies against IRF3 (PA5-20086), pIRF3 (Ser385) (PA5-38285), IRF7 (PA5-79519), pIRF7 (Ser477) (PA5-64834), GPX4 (PA5-79321), IκBα (PA5-22120), Complexes I (NDUFB8) (459210), III (UQCRC2) (PA5-30204) and IV (COX4) (A21348) were purchased from ThermoFisher. HMW (TLRL-PIC) and LMW Poly(I:C) (TLRL-PICW) were obtained from InvivoGen. *N*-acetylcysteine (NAC) (A9165), antimycin A (AA) (A8674), rotenone (ROT) (R8875), oligomycin (OM) (O4876), carbonyl cyanide-p-trifluoromethoxyphenylhydrazone (FCCP) (C2920), SQ1EL-1 (SML1948) and S3QEL-2 (SML1554) were acquired from Sigma-Aldrich. Genipin (sc-203057) was obtained from Santa Cruz. The IFN-α/IFN-β 2-Plex Mouse ProcartaPlex^TM^ Luminex Panel kit (EPX020-22187-901) was from Invitrogen. Antibodies detecting Complex II (SDHB) (ab14714) were from Abcam while antibodies recognizing SOD2 (13141S) were from Cell Signaling Technology.

### Animals

All animal procedures were approved by the Carleton University Animal Care Committee and conducted in accordance with Canadian Council for Animal Care guidelines. Bone marrow cells were isolated from the tibias and femurs of 6-13-week-old C57BL/6 male mice as previously described^67^ and cryopreserved in a 90% FBS/10% DMSO solution until use.

### Culture and Treatment of Bone Marrow-derived Macrophages (BMDMs)

As previously described^18,67^, mouse bone marrow progenitors were differentiated for 10 days in high glucose DMEM supplemented with 10% FBS, 1% PenStrep and 15% L929-conditioned medium on 100mm non-tissue culture treated Petri dishes. On day ten, BMDMs were harvested, counted, and seeded at 1×10^6^ cells/mL onto tissue culture or non-tissue culture treated plates in complete DMEM with 25mM (high) or 0.5mM (low) glucose, and rested overnight. Low glucose media was prepared by supplementing glucose/glutamine/pyruvate/phenol red-free DMEM with 0.5mM glucose, 4mM L-glutamine and 1mM sodium pyruvate before adding 10% FBS, 1% PenStrep and 15% L929-conditioned medium. BMDMs were then stimulated with HMW or LMW Poly(I:C) (1ng/mL-10µg/mL) for 1-18 hrs depending on the assay. For OXPHOS/mtROS inhibition studies, BMDMs were co-treated with 500μM MT, 5mM NAC, 5μM S1QEL, 5μM S3QEL-2 or 50μM genipin.

### Measurement of Cytokine Secretion

Culture supernatants were collected, centrifuged to remove cellular debris and analyzed for IFN- α and IFN-β levels *via* IFN-α/IFN-β 2-Plex Mouse ProcartaPlex^TM^ Luminex Panel according to manufacturer’s instructions.

### Western Blot Analysis

Cells were lysed in RIPA buffer (Thermo-Fisher; 89900) supplemented with HALT^TM^ Protease and Phosphatase Inhibitor Cocktail (ThermoFisher; 78441). Total sample protein was measured *via* DC Protein Assay (Bio-Rad; 5000112). Afterwards, 30µg of protein/sample were resolved on 12% TGX Stain-Free^TM^ FastCast^TM^ Acrylamide gels (Bio-Rad; 1610185), using Tris/Glycine/SDS buffer (formulated from Bio-Rad 161-0732), and imaged with the Stain-Free^TM^ program of ChemiDoc^TM^ XR system (Bio-Rad). Resolved proteins were transferred onto PVDF membranes using the Trans-Blot^®^ Turbo^TM^ Transfer System (Bio-Rad; 170-4272), blocked overnight in 5% non-fat dry milk (w/v) in Tris-buffered saline (TBS) with 0.1% Tween-20 (TBST), then incubated with primary antibodies overnight in 5% BSA (w/v) in TBST. Horseradish peroxidase-conjugated secondary antibodies and Clarity^TM^ Western ECL Substrate (Bio-Rad; 1705060) were used for detection. Band intensity was normalized to total protein per lane, as quantified by the Stain-Free application^68^. For IRF3 and IRF7, phosphorylated protein levels were presented as the ratio over total protein levels.

### Assessment of Mitochondrial Function using Flow Cytometry

BMDMs (2×10^6^ cells) were plated onto 60mm non-tissue culture coated Petri dishes and rested overnight before an 18-hour treatment with 10ng/mL HMW or LMW Poly(I:C). Cells were collected, washed, and stained with fluorescent probes per manufacturer’s instructions. Mitochondrial abundance was measured using 150nM MitoTracker^TM^ Green in serum-free DMEM. MMP was evaluated using 10nM TMRM in complete low glucose DMEM. mtROS levels were monitored using 2.5μM MitoSOX^TM^ Red in PBS. Cells were incubated with appropriate probes at 37°C for 30 minutes prior to PBS washing and subsequent flow cytometry analysis. For PI staining, treated BMDMs were incubated on ice with 4ug/mL PI in PBS for 15 minutes prior to subsequent flow analysis.

Single cell fluorescence was quantified using an Attune NxT Flow Cytometer (ThermoFisher), capturing ≥10,000 events per sample, and analyzed using FlowJo Software (v10.0.7). BMDMs were first gated based on size (via forward scatter (FSC) and side scatter (SSC)) before subsequently gating on the single cell population (*via* SSC-H vs. SSC-A scatter plot). Data was then characterized based on the fluorescent profiles of each probe. For TMRM, MitoSOX^TM^ Red and PI, subpopulations were categorized as either high or low fluorescence cells. TMRM data was presented as the median fluorescence intensity (MFI) of the “TMRM high” group normalized to their respective control sample (as a fold change (FC) value) as well as the proportion of “TMRM low” cells. MitoSOX^TM^ Red data was reported as FC of the percentage of “MitoSOX^TM^ Red high” cells relative to the control. PI data was reported as FC of the percentage of “PI High” subpopulation normalized to the respective control sample. For MitoTracker^TM^ Green, MFI of the total population was normalized to the control and presented as FC.

### Characterization of Energy Metabolism of BMDMs

BMDMs were plated at 50,000 cells/well onto Seahorse XFp miniplates (Agilent Technologies) and rested overnight before stimulation with various concentrations of HMW or LMW Poly(I:C) for 18 hours. Extracellular acidification rate (ECAR) and oxygen consumption rate (OCR) were evaluated using a XFp Flux Analyzer (Agilent Technologies). Mitochondrial function was assessed using the Seahorse XFp Cell Mito Stress Test Kit (Agilent Technologies), *via* successive injections of OM, FCCP, and ROT/AA. OCR data were used to calculate spare respiratory capacity percentage (SRC%), ATP-production coupled respiration and proton leak.

### Statistical Analyses

Data was analyzed using GraphPad Prism (v10.5.0). The number of biological replicates is indicated in the figure legends and graphs represent the mean +/- SEM of replicates. Where applicable, matched, ordinary one-way ANOVA with post-tests or a matched, ordinary two-way ANOVA with post tests were performed (*p < 0.05, **p < 0.01, ***p < 0.001, ****p < 0.0001).

## Results

### HMW Poly(I:C) is associated with higher levels of type I IFN production

It is increasingly recognized that media composition shapes intracellular metabolic responses *in vitro*^57^. Given HIF-driven inflammatory responses to HMW or LMW Poly(I:C) can be amplified under low glucose conditions^66^, we evaluated whether these ligands differentially produce type I IFNs in high and low glucose. In high glucose media (25mM), BMDMs stimulated with 10ng/mL HMW or LMW secrete IFN-α and IFN-β, with HMW producing more IFN-β compared to LMW (Figure 1A, B). Interestingly, both ligands produce higher levels of both under low glucose (0.5mM) conditions (HMW-IFNα FC: 5.935, p=0.008; IFN-β FC: 2.972, p=0.006 and LMW - IFNα FC: 2.081, p=0.036; IFN-β FC: 3.349, p=0.003), with HMW producing more than LMW (Figure 1A, B). As low glucose conditions more closely reflect levels in lung, skeletal muscle, and brain^69–73^, all subsequent experiments used low glucose media. Next, to determine if these differences were associated with differential antiviral signaling, we evaluated changes in phosphorylated IRF3 (pIRF3) and pIRF7 at early (1-4 hrs) and later time points (18 hrs). Both pIRF3 and pIRF7 were increased following TLR3 engagement (Figure 1C), which was sustained 18 hours post activation (Figure 1D), but did not account for differences in cytokine production between ligands.

**Figure 1:**
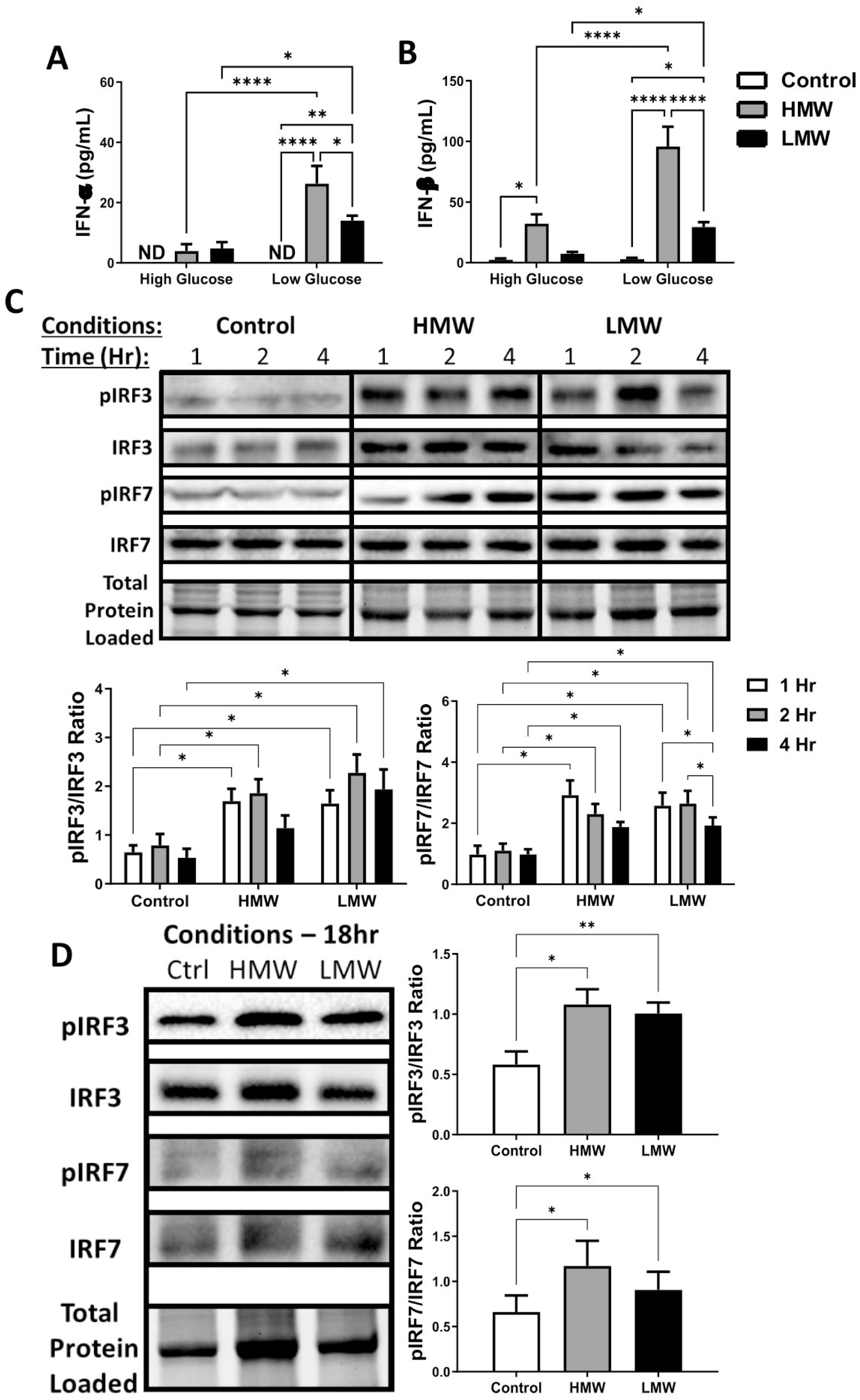
HMW Poly (I:C) engagement induced higher magnitude of Type I IFN signaling and cytokine production. BMDMs were treated with 10ng/mL HMW or 10ng/mL LMW Poly(I:C) for 18 hours in complete DMEM with 0.5mM glucose. Cytokine expression (IFN-α (**A**) and IFN-β (**B**)) was evaluated in culture supernatant. Total cell lysates were harvested to measure total and phosphorylated IRF3 and IRF7 (**C, D**) expression using immunoblotting at 1, 2 & 4 hours (**C**) as well as 18 hours (**D**) after treatment. Band intensity was normalized to the total protein levels in each of their respective lanes using the Bio-Rad Stain-Free Application. The samples shown in Figure 1c were analyzed using two separate blots imaged with the same exposure based on the same principles. Afterwards, the expression of pIRF3 and pIRF7 were normalized relative to the intensity of their total forms (IRF3 and IRF7) and presented as a ratio. Data represents mean ± SEM of four individual mice (*p < 0.05, **p < 0.01, ***p < 0.001, and ****p < 0.0001). ND = Not detected.

### HMW Poly(I:C) is associated with a more pronounced loss of SRC and MMP

Next, we assessed if HMW and LMW stimulation is associated with differential OXPHOS reprogramming using Agilent’s Cell Mito Stress Test kit and the Seahorse XFp Analyzer. At 10ng/mL, neither ligand altered basal respiration, ATP production or proton leak compared to controls (Figure 2A, C-D). However, both significantly decreased %SRC, with HMW having a more pronounced effect vs. LMW Poly(I:C) (Figure 2B). To rule out differences in ligand potency, we compared BMDMs stimulated with 10 ng/mL HMW or 100 ng/mL LMW Poly(I:C), which elicit comparable cytokine responses (Supplementary Figure 1A, B). At these concentrations, we observed a similar difference in OXPHOS function between the two ligands (Supplementary Figure 1C). Further at 10 µg/mL, which mimic responses to higher viral levels and can trigger a robust inflammatory response^18,74^, HMW and LMW Poly(I:C) similarly reduced basal respiration, %SRC, ATP production but differentially decreased proton leak, which was more pronounced with HMW (Supplementary Figure 1D). These findings suggest that TLR3 engagement with dsRNA of diverse lengths induce differential mitochondrial reprogramming. This aligns with recent crystal structures showing that longer dsRNAs form more highly organized lateral multimeric complexes^75–77^.

**Figure 2:**
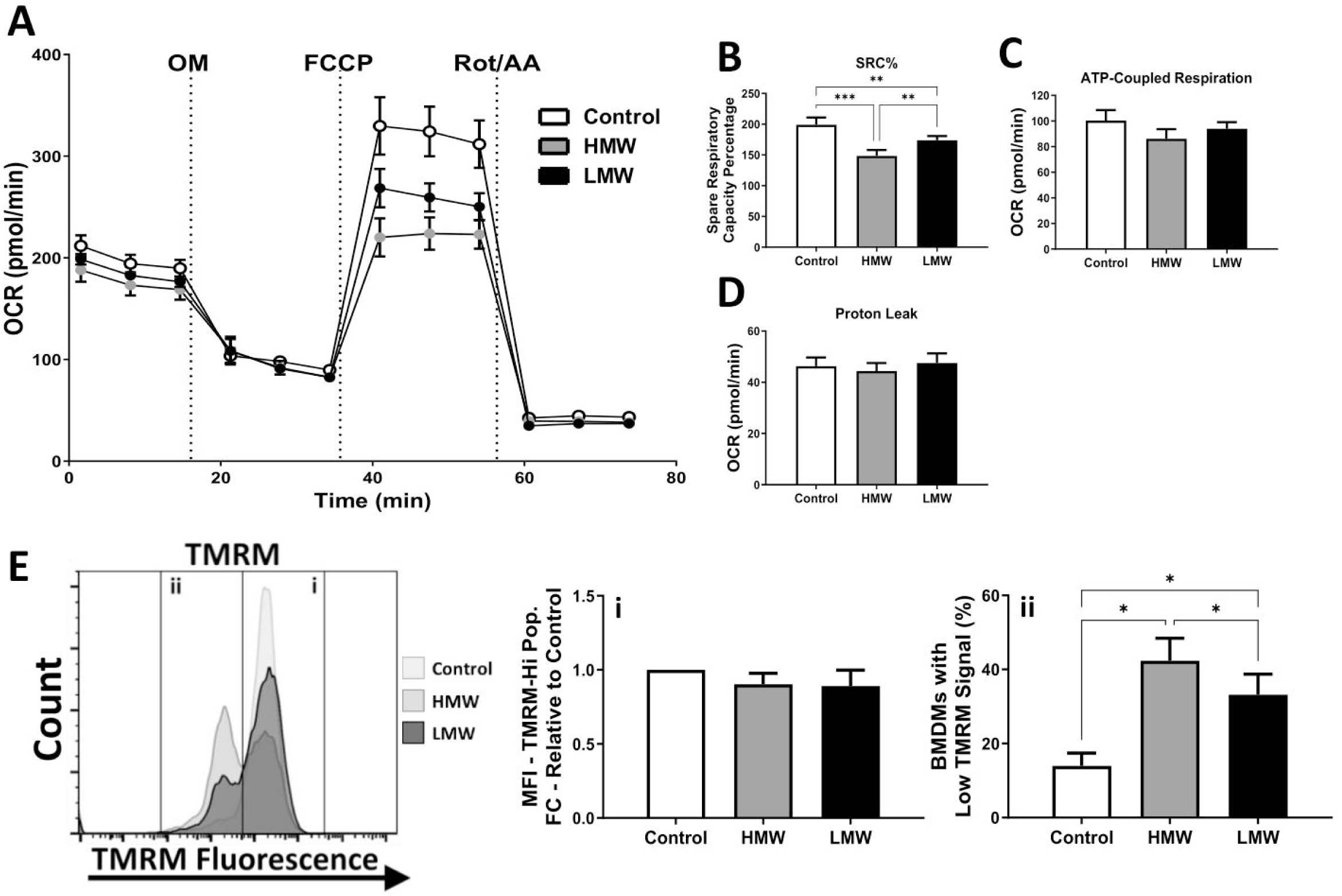
HMW and LMW Poly(I:C) have divergent effects on mitochondrial respiration and MMP. BMDMs were plated onto Seahorse XFp miniplates and subsequently stimulated with HMW or LMW for 18 hours (**A**) prior to assessing OXPHOS activity using the Cell Mito Stress Test kit (*via* consecutive Oligomycin (OM), Carbonyl cyanide-p-trifluoromethoxyphenylhydrazone (FCCP), and Rotenone (Rot)/Antimycin A (AA) injections). This allowed for the quantification of OXPHOS features such as spare respiratory capacity percentage (SRC%; **B**), ATP production (**C**) and proton leak (**D**). Tetramethylrhodamine (TMRM) staining was used to measure changes in mitochondrial membrane potential (**E**) and reported as the average MFI of the TMRM high population (**i**) as well as the percentage of BMDM with low MMP (**ii**). Data represents mean ± SEM of four individual mice. (*p < 0.05, **p < 0.01, ***p < 0.001, and ****p < 0.0001).

Since MMP is directly influenced by mitochondrial respiration^78^, we evaluated if HMW/LMW stimulation was associated with changes in MMP using the fluorescent dye TMRM. We found that both ligands increased the proportion of cells with low TMRM sequestration and that this increase was more pronounced during HMW activation (p=0.0422) (Figure 2E). Importantly, these changes were not associated with any change in mitochondrial content (Supplementary Figure 2). Overall, these findings indicate that while TLR3 engagement preserves OXPHOS activity, a fraction of activated BMDMs display low or undetectable MMP, which is influenced by dsRNA length.

### Poly(I:C)-driven type I IFN responses is dependent on UCP2-mediated proton leak

The UCP family are critical inner mitochondrial membrane proteins that regulate uncoupling proton movement from mitochondrial ATP production while dissipating MMP^79^. UCP2 is unique because it transports C4 metabolites (e.g., malate, oxaloacetate, and α-ketoglutarate) in exchange for protons^79^. Given UCP2 has been shown to modulate inflammatory responses^80–84^, we assessed whether the differences in MMP between ligands were linked to its altered expression. Using western blot, we found that both HMW and LMW activation similarly induced UCP2 expression (Figure 3A). We then used the UCP2 inhibitor genipin^80,85^ to investigate how UCP2-driven proton leak contributes to Poly(I:C) activation. Consistent with LPS^86^, UCP2 inhibition resulted in elevated MMP and significantly reduced the proportion of low TMRM cells (Figure 3B, C). This was accompanied by a severe reduction in IFN-β secretion, irrespective of ligand length (HMW: ↓77.55%, p<0.0001; LMW: ↓40.20%, p<0.0001) (Figure 3C). Interestingly, both ligands showed similar reductions inIFN-β, suggesting that disrupted active proton movement impairs TLR3 ligand-independent antiviral responses but not in a ligand specific manner.

**Figure 3:**
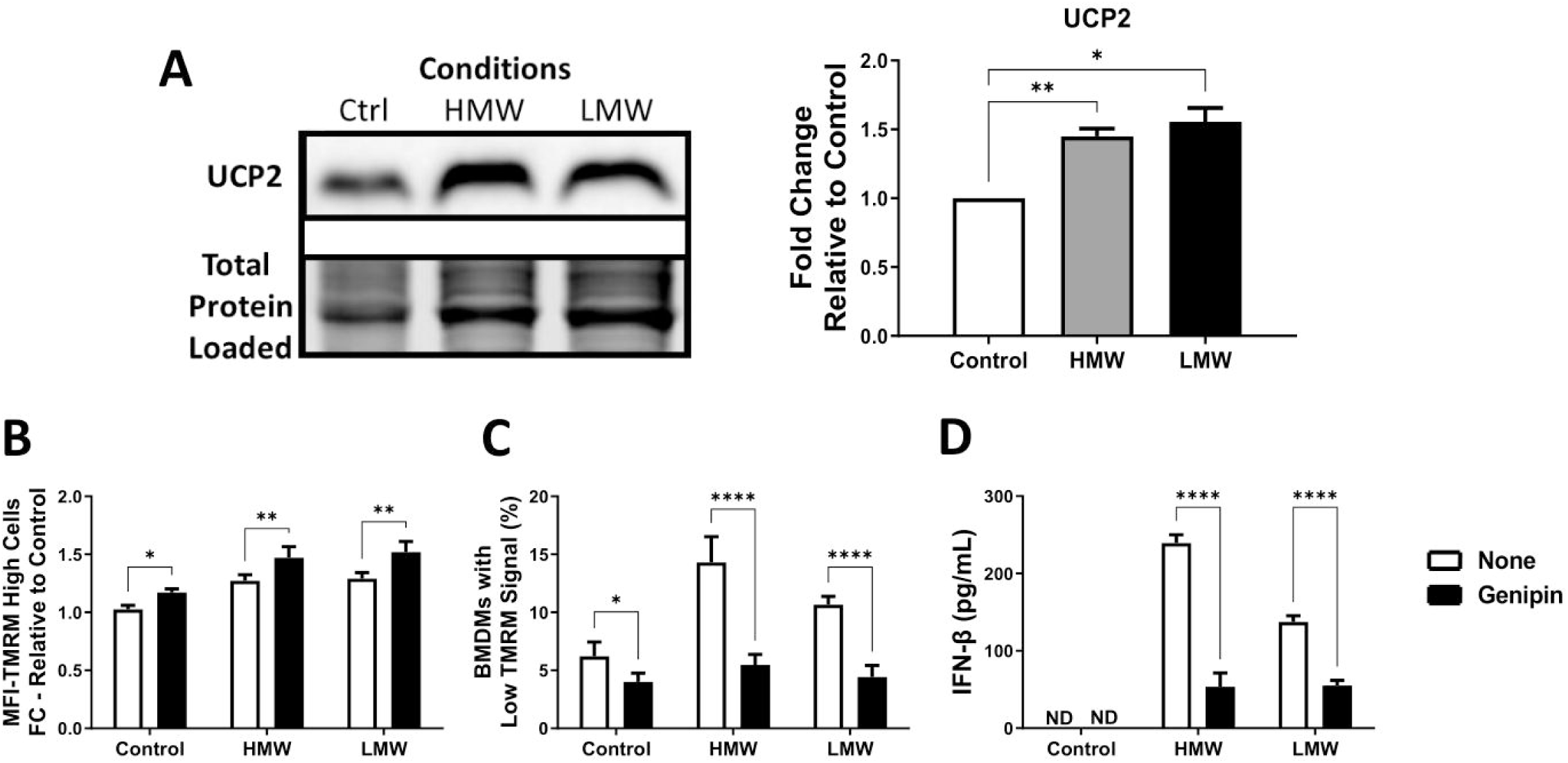
Genipin inhibition of UCP2-mediated proton leak results in severe but equivalent loss of IFN-β production during HMW and LMW activation. UCP2 protein expression in HMW- and LMW-stimulated BMDMs was quantified *via* immunoblotting (**A**). Band intensity was normalized to the total protein levels in each of their respective lanes using the Bio-Rad Stain-Free Application. All samples on each blot in this figure, were loaded onto a single blot. Afterwards, the expression of each protein was normalized to the control and presented as a fold change value. The importance of UCP2-mediated proton leak were assessed using genipin. After 18 hours of co-treatment, TMRM staining was done to measure the average MFI of the TMRM high population (**B**) as well as the percentage of BMDM with low MMP (**C**). Cell supernatant was collected after treatment and IFN-β secretion was quantified via ELISA (**D**). Data represents mean ± SEM of four individual mice (*p < 0.05, **p < 0.01, ***p < 0.001, and ****p < 0.0001). ND = Not detected.

### Differential Modulation of Complex I Expression between HMW vs. LMW Poly(I:C) drives differences in mitochondrial ROS accumulation

To better understand the drivers of altered mitochondrial function, we next evaluated changes in the proton-pumping ETC complexes (complexes I, III and IV) *via* immunoblotting. Following stimulation, both HMW and LMW Poly(I:C) decreased Complex I (NDUFB8; HMW FC: 0.664, LMW FC: 0.524) and Complex IV expression (COX4; HMW FC: 0.767, LMW FC: 0.751) while increasing Complex III expression (UQCRC2; HMW FC: 2.224, LMW FC: 2.348) (Figure 4). Notably, Complex I downregulation was more pronounced in LMW vs. HMW Poly(I:C) (p=0.035) suggesting it may play an important role in HMW vs. LMW mediated responses.

**Figure 4:**
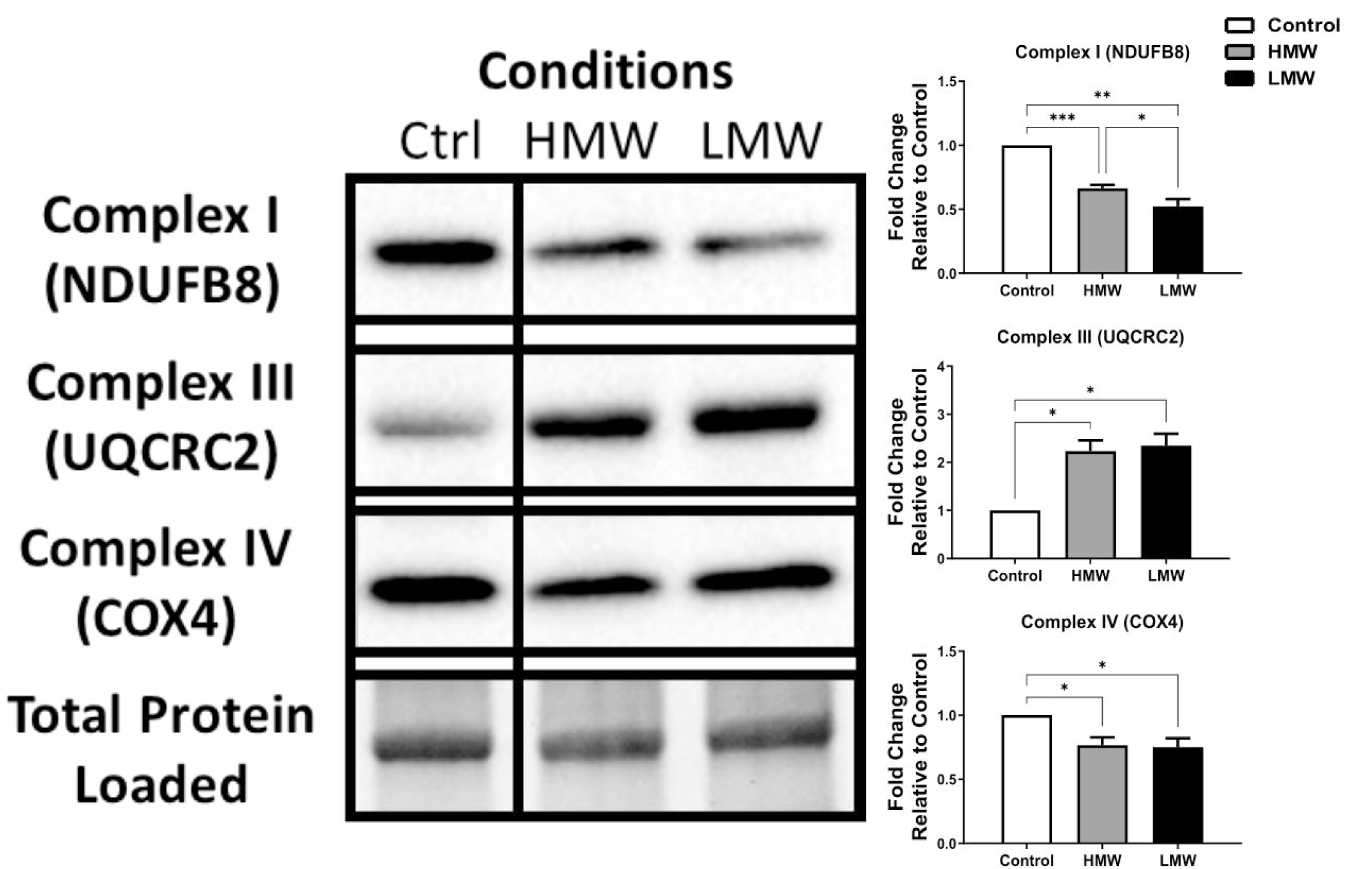
dsRNA length-dependent TLR3 engagement determines the degree of Complex I expression. Macrophages treated with HMW or LMW for 18 hours were evaluated for differences in mitochondrial function. Protein levels of complexes I, III and IV of the ETC was quantified *via* immunoblotting. Band intensity was normalized to the total protein levels in each of their respective lanes using the Bio-Rad Stain-Free Application. All samples on each blot in this figure, were loaded onto a single blot but cropped to focus on the shown samples. Afterwards, the expression of each protein was normalized to the control and presented as a fold change value. Data represents mean ± SEM of five individual mice (*p < 0.05, **p < 0.01, ***p < 0.001, and ****p < 0.0001).

Given these changes in ETC architecture, we next investigated if they resulted in differential mtROS accumulation using MitoSOX^TM^ Red to measure mitochondrial superoxide production. As expected^18^, TLR3 engagement increased mitochondrial superoxide production, which was more pronounced following HMW engagement (p=0.0067) (Figure 5A). This accumulation occurred without any upregulation in SOD2 or mtGPX4 expression (Figure 5B, C), suggesting the differences in mtROS accumulation are not linked to their respective antioxidant responses. To determine if mtROS contributes to antiviral signaling, BMDMs were co-treated with two antioxidants (MitoTEMPO [MT], N-acetylcysteine [NAC]), neither of which significantly affect cell viability (Supplementary Figure 3). MT scavenges superoxide from active mitochondria whereas NAC targets total cellular ROS production, yet both nearly eliminated any type I IFN induction following TLR3 engagement (Figure 5D, E) demonstrating the importance of mtROS to TLR3-mediated antiviral responses.

**Figure 5:**
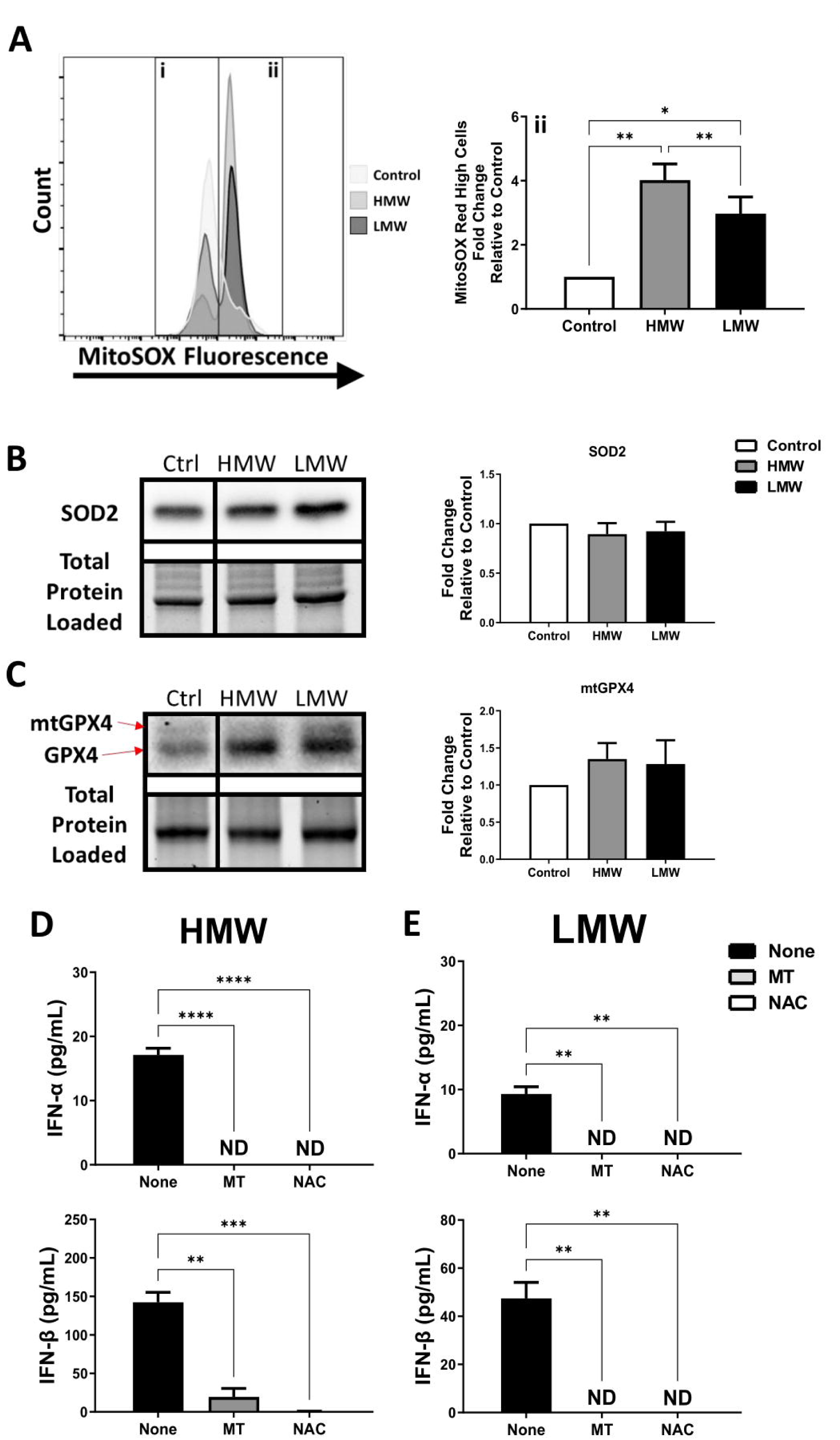
Scavenging mtROS blunts Poly(I:C)-mediated type I IFN production. HMW- and LMW-stimulated BMDMs were probed for changes in redox metabolism. Mitochondrial superoxide levels were measured using MitoSOX^TM^ Red (**A**) and reported as the percentage of BMDM with high MitoSOX Red signal (**ii**). Expression levels of antioxidant proteins superoxide dismutase 2 (SOD2) (**B**) and mitochondrial glutathione peroxidase 4 (mtGPX4) (**C**) were quantified using immunoblotting. Band intensity was normalized to the total protein levels in each of their respective lanes using the Bio-Rad Stain-Free Application. All samples on each blot in this figure, were loaded onto a single blot. Afterwards, the expression of SOD2 and mtGPX4 was normalized to the control and presented as a fold change value. HMW- (**D**) or LMW-activated BMDMs (**E**) were also co-treated with two different ROS scavengers (MitoTEMPO [MT], N-acetylcysteine [NAC]) to better understand the importance of ROS to drive antiviral responses. IFN-α and IFN-β production were quantified after 18 hours *via* ELISA. Data represents mean ± SEM of four individual mice (*p < 0.05, **p < 0.01, ***p < 0.001, and ****p < 0.0001). ND = Not detected.

### Mitochondrial ROS from Complexes I and III differentially drive type I IFN responses following stimulation with HMW or LMW Poly(I:C)

Since electron leakage at complexes I and III generates mitochondrial superoxide^87,88^, we assessed whether preventing mtROS production while preserving OXPHOS, using S1QEL (Complex I) or S3QEL (Complex III)^89,90^, differentially alters HMW/LMW IFN productions. While these inhibitors did not significantly affect cell viability (Supplementary Figure 3), they had distinct effects on HMW vs. LMW Poly(I:C) responses. We found that LMW responses required mtROS from both complexes to drive type I IFN production (Figure 6B), whereas HMW-induced IFN-β did not depend on Complex I-derived mtROS (Figure 6A). In fact, S1QEL inhibition increased IFN-β production (↑27%, p=0.0386), which was accompanied by similar pIRF3 levels in HMW-treated cells (Figure 6C). Given that IRF3 activation drives early type I IFN responses and IRF7 phosphorylation support late phase antiviral responses, including IFN-α and further IFN-β production^91^, these data suggest that Complex I-derived mtROS may suppress early type I IFN responses during HMW stimulation. Furthermore, our work implies that targeting Complex I may be a novel approach for augmenting antiviral therapies.

**Figure 6:**
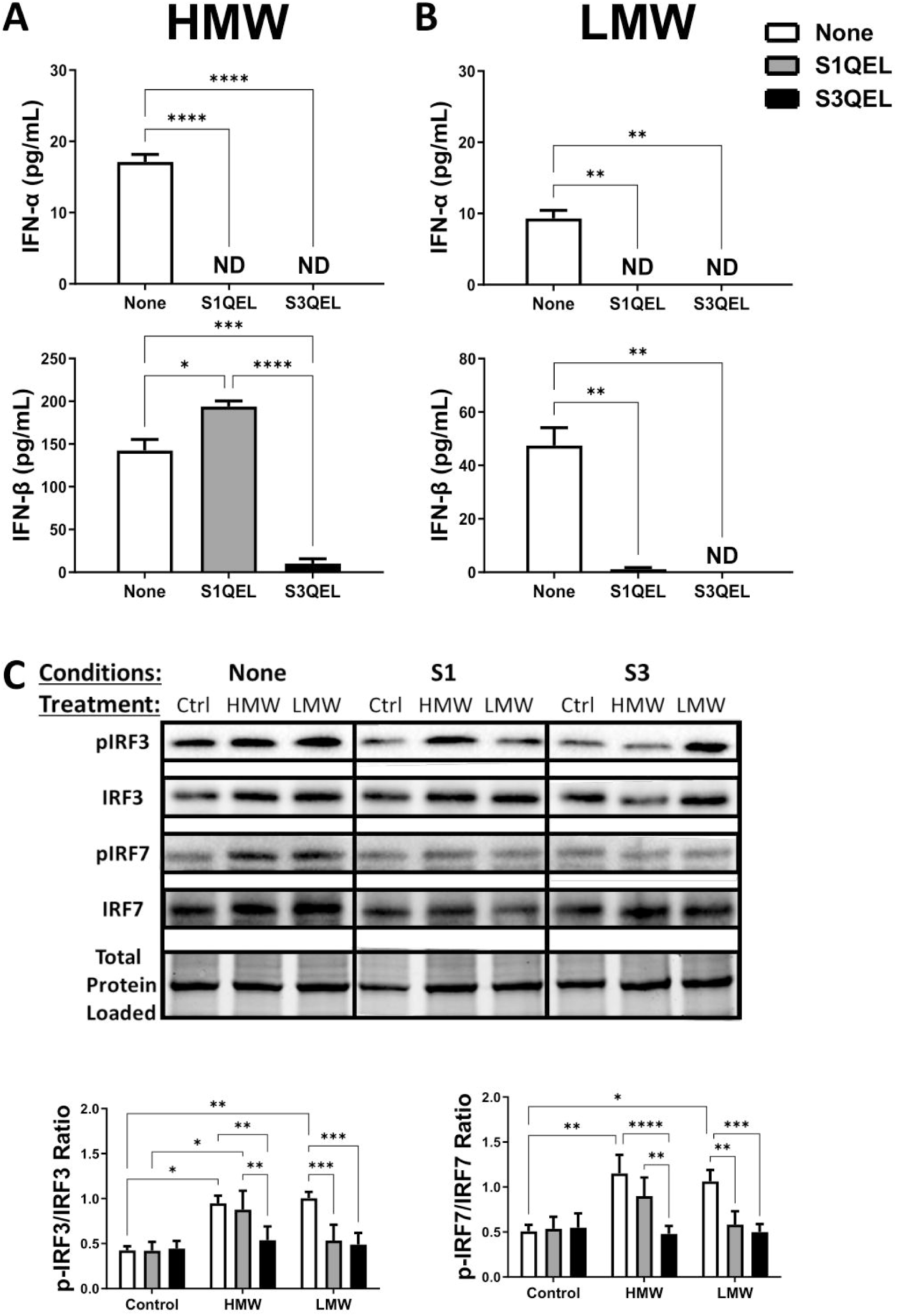
Preventing the production of Complex I-generated mtROS can unlock HMW-mediated Type I IFN production *via* increased IRF3/7 activation BMDMs treated with either HMW. (**A**) or LMW (**B**) were co-treated with specific suppressors of electron leakage from Complex I (S1QEL) or Complex III (S3QEL) to better understand the importance of mtROS-producing sites to driving antiviral response. IFN-α and IFN-β cytokine production were quantified after 18 hours. Total cell lysates were harvested to measure total and phosphorylated IRF3 (**C**) and IRF7 (**D**) after S1QEL and S3QEL co-treatment using immunoblotting. Band intensity was normalized to the total protein levels in each of their respective lanes using the Bio-Rad Stain-Free Application. All samples on each blot in this figure, were loaded onto a single blot. Afterwards, the expression of IRF3 and IRF7 was normalized relative to the intensity of their phosphorylated forms (pIRF3 and pIRF7) presented as a ratio of phosphorylation signal. Data represents mean ± SEM of four individual mice (*p < 0.05, **p < 0.01, ***p < 0.001, and ****p < 0.0001). ND = Not detected.

## Discussion

Innate sensing of PAMPs by PRRs, including dsRNA/TLR3, is critical to mounting an effective effector response. dsRNA is found in viruses capable of infecting everything from bacteria to animals^59,61,62,92^. These viral genomes can be monopartite (e.g. L-A virus) or segmented (e.g. *Megabirnaviridae*: 2 segments [7–9kb]; *Partitiviridae*: 2–3 segments [1.4–2.3kb]; *Reoviridae*: 10– 12 segments [0.7–5kb])^59–62^. Emerging evidence suggests that endogenous dsRNA^63,93,94^ and mRNA^95,96^, released by infected or dying cells, also engage TLR3 signaling and thus requires detection of diverse RNA lengths/structures (>40-50 bp)^58,92^. Poly(I:C), a synthetic dsRNA of variable lengths, is widely used as a viral mimic to assess antiviral reponses^97–99^ as well as in immunotherapies (e.g. adjuvants for viral and cancer vaccines)^100–104^. Yet, dsRNA length influences TLR3 affinity and clustering, driving distinct activation states across cell types^64,65,75–77,105^.More insight is required to optimize their potential in immunotherapies.

In this study, we found that TLR3 engagement by long (HMW) vs. short (LMW) dsRNA differentially reprograms mitochondrial function to support early ligand-specific antiviral responses. These ligands induced differential mtROS accumulation that drove IRF3/7 phosphorylation and type I IFN production. Although OXPHOS preservation and UCP2-facilitated proton leak were required for TLR3 signaling, they did not differentiate HMW and LMW responses. Instead, differences in Complex I expression and mtROS production separated the two activation states. Importantly, Complex I acted as a rheostat whose mtROS suppresses type I IFN secretion and limiting it can boost HMW-driven antiviral responses, identifying a novel mechanism to selectively induce antiviral response in macrophages.

Mitochondrial function and antiviral responses are intimately linked, with sustained OXPHOS activity and MMP key for effective signaling^16,19–24^. Studies have shown that RIG-I-like receptor (RLR)-mediated macrophage responses require OXPHOS activity as targeting MMP or ETC electron flow increased viral susceptibility^19,43^. In contrast, RIG-I activation reduces glycolysis by disrupting hexokinase 2 (HK2)-MAVS mitochondrial interactions while redirecting glucose towards the pentose phosphate pathway and hexosamine biosynthesis to support type III and type I IFN production, respectively^41,47^. Unlike LPS-induced inflammatory reprogramming^15,16,22,29^, we found that TLR3-mediated responses, irrespective of dsRNA length, is facilitated by OXPHOS activity. Accordingly, Albers and colleagues^106^ showed that knocking out IRF5, a driver of inflammatory (e.g. *Il6, Il12, Tnf*)^107,108^, glycolytic (e.g*. Hk2*) and mitochondrial gene (e.g. *Idh2*) expression, diminished OCR of Poly(I:C)-stimulated BMDMs. Moreover, increased OXPHOS in TLR9-activated plasmacytoid dendritic cells is driven by an autocrine type I IFN-mediated boost in fatty acid oxidation^109^, but this remains to be explored in TLR3 responses.

MMP and ultimately OXPHOS are shaped by proton leak, uncoupling proton transport from OXPHOS and ATP production^78^. We found that TLR3-mediated antiviral responses was influenced by UCP2-driven active proton leak. This proton leak likely contributes to the increased proportion of low MMP BMDMs largely responsible for type I IFN secretion. In contrast, UCP2 overexpression in mouse embryonic fibroblasts enhanced active proton leak but inactivated MAVS-mediated RIG-I signaling^43^. There has also been some debate regarding the effect of proton leak on mtROS production. Koshiba et al.^43^ showed that UCP2 overexpression during RIG-I signaling attenuated mtROS generation which dampens the antiviral response^39^. Whereas we found that UCP2-driven proton leak was associated with mtROS production during TLR3 engagement, enabling IRF3/7 activation. Despite these differences, both RIG-I- and TLR3-mediated antiviral signaling is driven in part by mtROS^18,39,110^.

Previous work has shown that complex III-mediated mtROS facilitates effector function by driving macrophage pro-inflammatory^40^, anti-inflammatory^111^ and antiviral signalling^18^. We further explored ETC function in macrophages, demonstrating that Complex I also regulates type I IFN responses following HMW and LMW Poly(I:C) stimulation. Unlike complex III, complex I expression is downregulated, more markedly with LMW. We speculate that this downregulation accounts for the mtROS differences to protect against mtROS accumulation and subsequent damage. Further, reduced Complex I expression in LMW-stimulated BMDMs was associated with more cells with detectable MMP and a higher SRC% relative to HMW-stimulated BMDMs. Both parameters suggest decreased mitochondrial dysfunction in LMW vs. HMW activation, especially as low SRC is indicative of mitochondrial impairment^112,113^. Consistent with this, HMW activation supports the highest mtROS induction while preventing complex I-derived mtROS increased IFN-β production. This is possible that preventing electron leakage at complex I using S1QEL may force electrons to travel towards complex III, where the mtROS generated would promote antiviral signaling. This may be due to a dependency of early antiviral immune responses on OXPHOS and ETC supercomplex dynamics, which optimize electron transfer to reduce mtROS generation^114,115^.

While antibacterial responses involve supercomplex disassembly to increase mtROS and drive inflammasome activation^21,51^, information regarding their role in viral infections remains limited. Champagne et al.^116^ shown that methylation-controlled J protein deficiency enhances CD8+ T cell-driven Influenza immunity *via* increased supercomplex assembly. Yet Bladanta et al.^117^ demonstrated that interferon-stimulating gene ISG15 KO macrophages reduced free Complex I and boosted supercomplex assembly but heightened vaccinia susceptibility^117^. Recent work identified components of Complex IV participate in a conserved mitochondrial stress response (MISTR) that alters ETC/supercomplex dynamics during inflammatory/antiviral/hypoxic responses^118–121^. The IFN-induced MISTRAV/C15orf48 can facilitate mitochondrial fragmentation, mtROS/mtDNA release to drive STING-mediated tumour-associated macrophage activation^122^. Further studies are needed to determine how ETC supercomplex dynamics influence macrophage antiviral responses.

In summary, we show that TLR3 engagement alters ETC complex architecture based on dsRNA length, both requiring OXPHOS to mount mtROS-driven antiviral responses. Yet, both ligands differ in Complex I expression, causing differential respiratory capacity, MMP and mtROS production. Interestingly, blocking complex I superoxide production without impeding electron flow significantly increased HMW-derived antiviral signaling. These findings highlight macrophage functional plasticity and demonstrate how ETC dynamics can fine-tune antiviral responses, offering potential strategies to optimize macrophage-targeted therapies.

## Acknowledgements

This manuscript was supported by grant RGPIN-2019-06214 from the Natural Sciences and Engineering Research Council of Canada.

## Authorship Contribution Statement

DA and EC planned all the experiments, while DA, MD, AH, OA, DR, MES, ZV and RM conducted the experiments under the supervision of AG and EC. DA, AJ, AE, and AA harvested the mouse bone marrow progenitor cells used in this study. DA, MD, AH, DR, MES, ZV, AK, AG and EC contributed to the curation, analysis and interpreting of the data. All authors were involved in original draft of this manuscript as well as the review, editing and revision of the manuscript.

