## Supplemental Figures 1-3 for "Nucleic acid strand length governs mitochondrial reprogramming and mtROS-associated antiviral responses following TLR3 engagement"

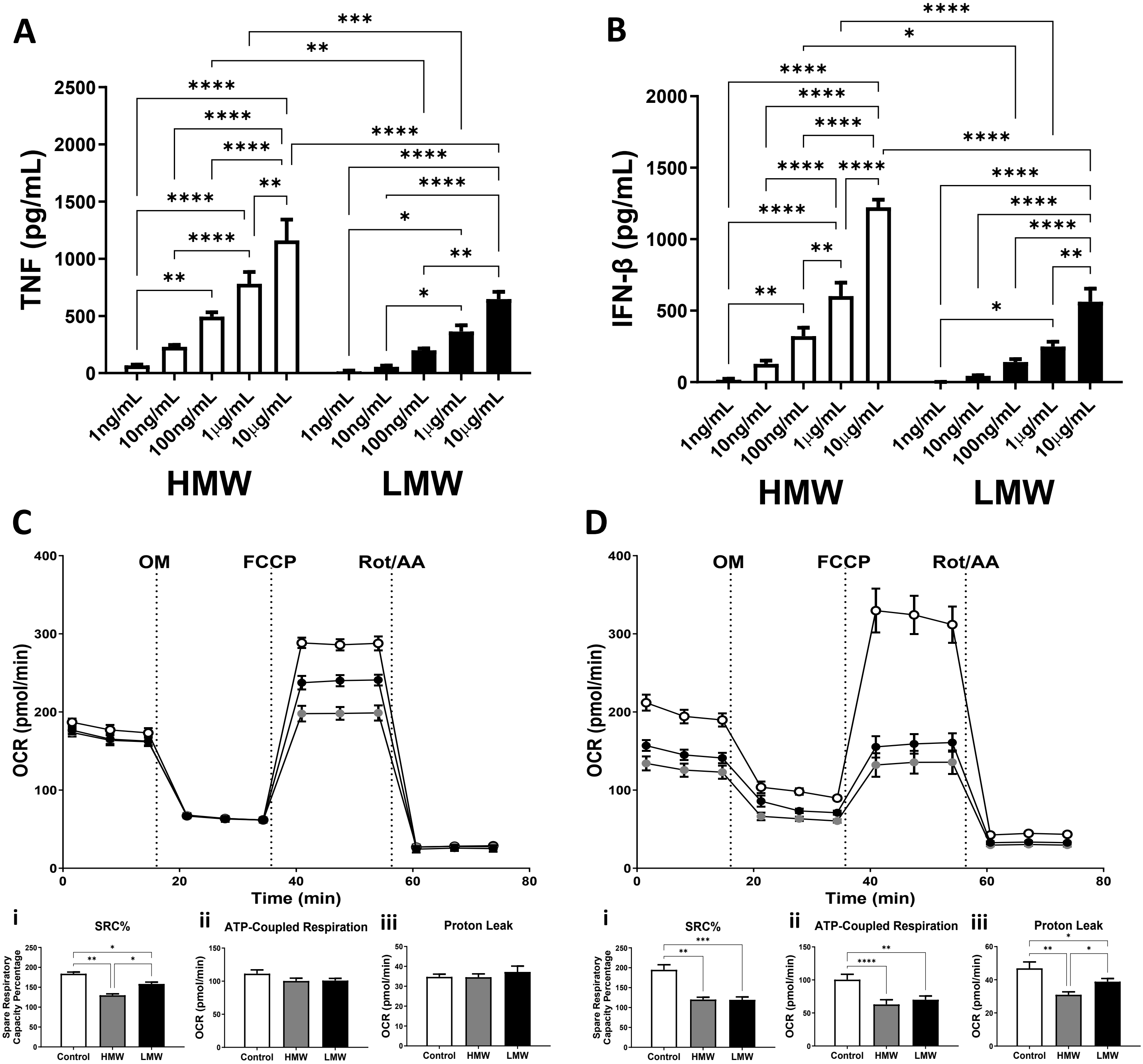


**Supplementary Figure 1: Comparable levels of HMW vs LMW activation does not equate to similar mitochondrial reprogramming.** BMDMs were stimulated with a range of HMW or LMW concentrations (1ng/mL to 10μg/mL) for 18 hours in low glucose media and subsequently assessed for their respective inductions of TNF (**A**) and IFN-β (**B**). Based on similar TNF/IFN-β production (highlighted in red boxes), BMDMs were seeded onto Seahorse XFp miniplates and treated with either 10 ng/mL HMW or 100ng/mL LMW for 18 hours to assess differences in mitochondrial reprogramming (**C**). Similarly, seeded BMDMs were also treated with 10ug/mL of either HMW or LMW for 18 hours (**D**). The treated cells were assessed for changes in OXPHOS activity using the Cell Mito Stress Test kit and subsequently assessed for OXPHOS features such as SRC% (**i**), ATP production (**ii**), and proton leak (**iii**). Data represents mean ± SEM of four (**A, B, D**) or three (**C**) individual mice. (*$p < 0.05$, **$p < 0.01$, ***$p < 0.001$, and ****$p < 0.0001$).

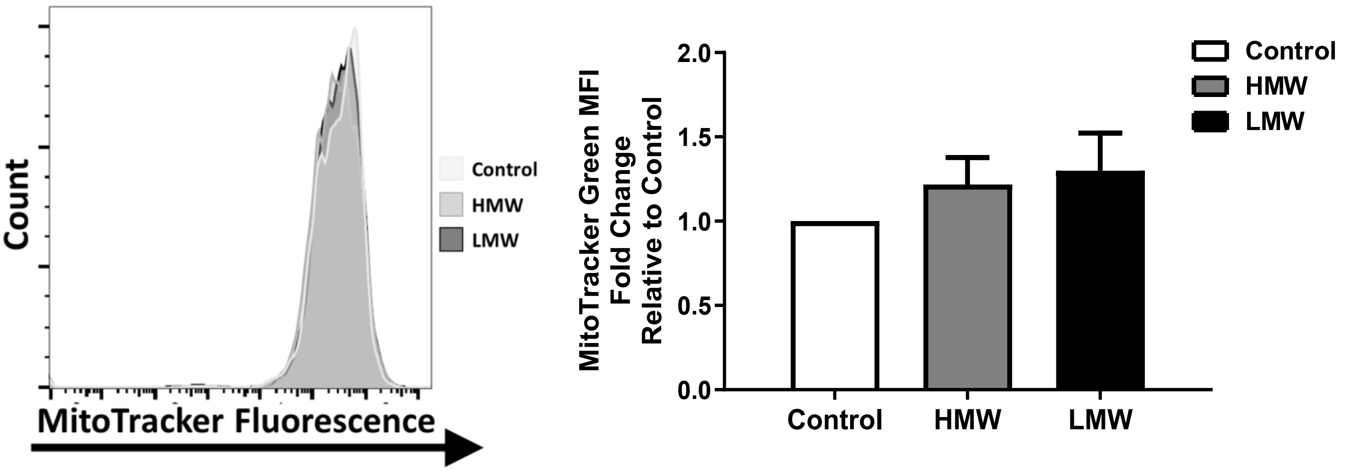


**Supplementary Figure 2: TLR3 engagement does not alter mitochondrial mass.** BMDMs treated with HMW or LMW for 18 hours in high or low glucose media were evaluated for changes in mitochondrial abundance. MitoTracker Green staining was used to measure mitochondrial abundance. Data represents mean ± SEM of four individual mice (*p < 0.05, **p < 0.01, ***p < 0.001, and ****p < 0.0001).

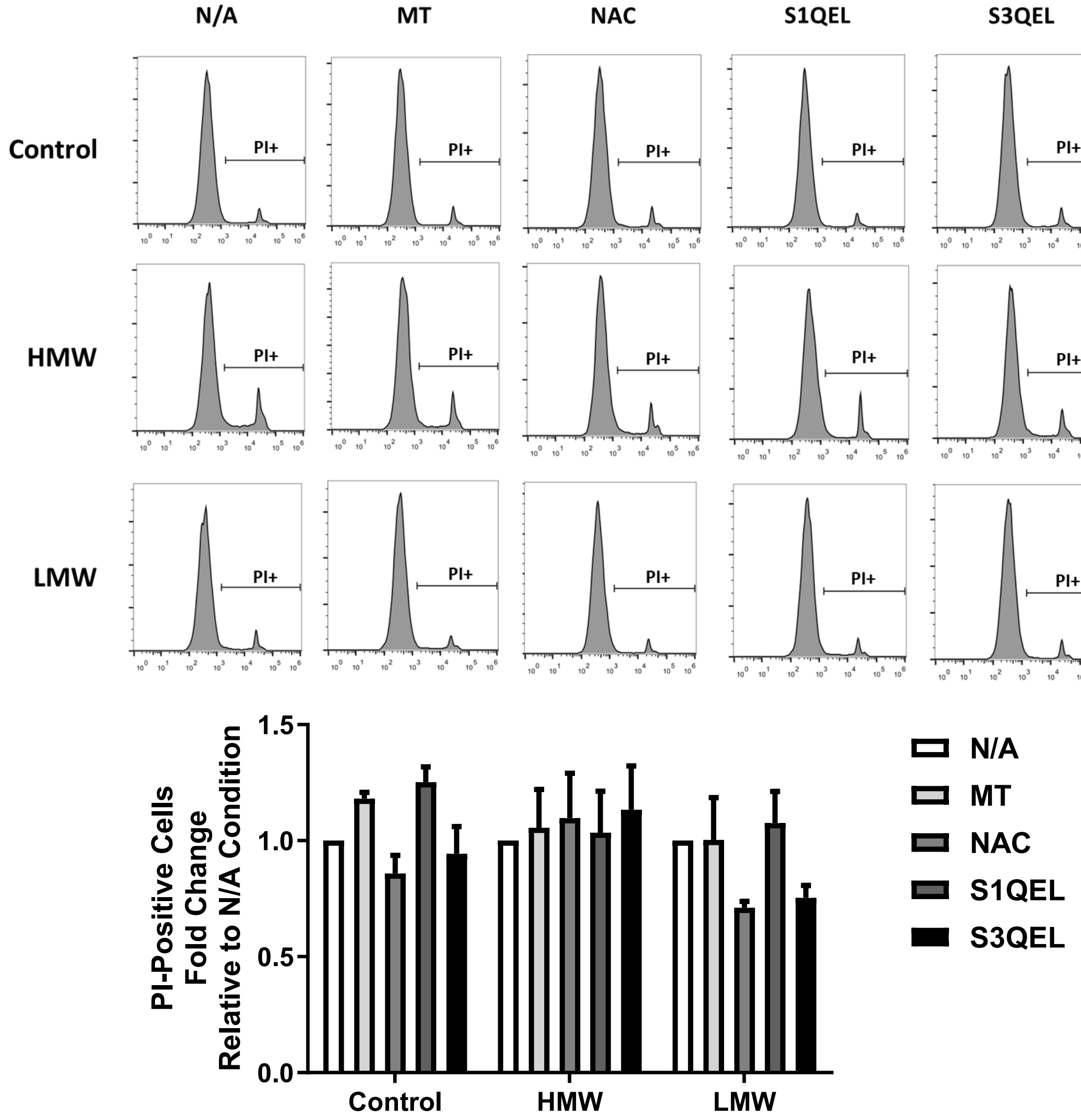


**Supplementary Figure 3: ROS inhibitors and TLR3 ligand co-treatment does not reduce cell viability.** BMDMs were co-treated with HMW or LMW and a panel of ROS inhibitors for 18 hours in low glucose media and subsequently assessed for differences in cell death using Propidium Iodide (PI). PI staining data was presented as fold change of PI-positive cells relative to each N/A condition. Data represents mean ± SEM of three individual mice (*p < 0.05, **p < 0.01, ***p < 0.001, and ****p < 0.0001).
